# Effects of C3aR activated mast cells in eosinophilic esophagitis

**DOI:** 10.1101/2025.05.23.655738

**Authors:** Simin Zhang, Dalila Cavallaro, Ming Xia, Netali Ben-Baruch Morgenstern, Tetsuo Shoda, Garrett A Osswald, Scott Bolton, Vince Mukkada, Tahereh M. Derakhshan, Julia Dunn, Jennifer Felton, Mark Rochman, Julie M. Caldwell, Marc E. Rothenberg

**Affiliations:** Division of Rheumatology, Allergy and Immunology, Department of Medicine, University of Cincinnati; Division of Allergy and Immunology, Cincinnati Children’s Hospital Medical Center; Division of Gastroenterology, Hepatology, and Nutrition, Cincinnati Children’s Hospital Medical Center; Division of Rheumatology, Immunology and Allergy, Brigham and Women’s Hospital; Division of Gastroenterology, Hepatology and Nutrition, University of Colorado

## Abstract

**Background:** Eosinophilic esophagitis is a chronic food induced allergic inflammatory disease of the esophagus. Eosinophil-depleting antibodies have not shown significant improvement in clinical symptoms, which has turned attention to mast cells. These cells also accumulate in the esophagus of EoE at levels that correlate with symptoms, demonstrate extensive activation including degranulation, correlate and remain elevated and poised for reactivation even in EoE patients in remission as defined by reduced eosinophil levels. Herein, we examine a mechanism of mast cell activation in EoE.

**Methods:** Esophageal mast cell degranulation was quantitated by CD63 expression by flow cytometry. Mast cell degranulation activity of esophageal biopsies was examined employing a bioassay with human CD34 progenitor cell derived primary human mast cells. Bulk and single cell RNA sequencing data of the esophagus were analyzed for complement dysregulation and mast cell properties as a function of disease state. Human esophageal biopsies were stained for C3 and analyzed for C3a and C3 protein content. Primary human mast cells and fibroblasts were further interrogated by flow cytometry, immunostaining, and RNA sequencing.

**Results:** Esophageal mast cells demonstrated increased CD63 expression, a measure of degranulation, in subjects with active compared with remission EoE. Esophageal biopsy lysates induced mast cell degranulation. Analysis of esophageal bulk RNA sequencing data demonstrated that the mast cell marker *CPA3* strongly correlated with *C3AR1*. Accordingly, the level of C3a receptor on esophageal mast cells was proportional with CD63 expression. *C3AR1* expression was increased in the fibrostenotic endoscopic phenotype compared to normal appearing active EoE patients. Esophageal biopsies were notably enriched in the expression of multiple complement genes (including *C3AR1*, *C3*, *CFB, C1QA, C1QB*, and *C1QC*) some of which were increased in active vs remission EoE (*C3AR1*, *CFB, C1QA, C1QB*, and *C1QC*). Single cell RNA (scRNA) sequencing data revealed enrichment of *C3* in fibroblasts at levels higher in EoE compared with non-EoE controls. The ratio of C3a/C3 protein levels in the esophagus was increased in EoE compared to control. Mast cells and fibroblasts have increased proximity in EoE compared with non-EoE control subjects. Additionally, C3a stimulated mast cells to express increased CD117 and release multiple EoE-relevant cytokines including fibroblast active mediators including granzyme B, which is enriched in esophageal mast cells.

**Conclusion:** Taken together, our findings present evidence for the interplay of mast cells and fibroblasts via C3a in the pathogenesis of EoE.

## Introduction

Eosinophilic esophagitis (EoE) is a chronic, food-induced allergic inflammatory disease with an increasing prevalence, now representing the most common reason for chronic dysphagia in young adults^1, 2^. Despite eosinophils being pathognomonic to disease diagnosis, recent clinical trials with eosinophil-depleting antibodies have indicated that eosinophils are not the primary causal cell^3–5^; this has turned attention to other immune cells which may be driving disease. Esophageal mast cells (MCs) in patients with EoE have been shown to be activated as assessed by single-cell RNA sequencing^6^ and immunostaining^7^, but the mode of activation is unclear. Additionally, to date, the complement system has been understudied in EoE, but more recent data has suggested complement dysregulation^8^.

## Methods

### Patients

All human studies were approved by the Cincinnati Children’s Hospital Medical Center (CCHMC) institutional review board (IRB), and all human subjects provided written consent. Distal esophageal biopsy was used throughout the study because this is typically obtained during endoscopy, represents the conventional location of biopsies, and allows ready comparison to the previous transcriptomic studies,^9, 10^ which are generally limited to this region. Control patients were those who had gastrointestinal symptoms without a prior history of EoE who underwent esophageal biopsy and were found to have a peak eosinophil count ≤10 eosinophil per high powered field (eos/hpf). Remission EoE was defined as patients with a history of EoE diagnosis but with a current biopsy peak eosinophil count <15 eos/hpf. Active EoE was defined as patients with an esophageal biopsy peak eosinophil count ≥15 eos/hpf. Patients on an oral or swallowed steroid were excluded. Human esophageal biopsies were randomly selected from the Cincinnati Center for Eosinophilic Disorder biorepository without consideration of sex, race or atopic status. Patients on oral steroids were excluded. The tissues obtained for research were then placed in RNAlater buffer (Qiagen) and later processed for RNA extraction or lysed for protein analysis/bioassays.

### Flow cytometry

Fresh esophageal biopsies were processed as previously described^6^. Briefly, they were collected and treated with EDTA stripping buffer, and digested with collagenase. The digested tissue was passed through a 19G needle and filtered through a 70μm filter to make a single cell suspension. Cells were stained for viability using Blue Live/Dead stain and blocked with Fc block. Cells were stained with a panel of antibodies: CD45 (FITC), C3aR (PE), FcεRI (PE Cy7), CD117 (BV711), and CD63 (AF647). After the incubation, cells were fixed with paraformaldehyde. Flow cytometry was performed on LSR Fortessa (BD Biosciences, New Jersey, USA) and data were analyzed with FlowJo v10.6 (FlowJo LLC, Ashland, OR, USA).

### Culture of primary mast cells

Mobilized human peripheral blood CD34+ cells (1×10⁶), obtained from Cincinnati Children’s Hospital Medical Center Cell Processing Facility, were cultured in StemSpan™ SFEM medium (StemCell Technologies, Vancouver, CA) for seven weeks. The medium was supplemented with 100 ng/mL Stem Cell Factor (Peprotech, New Jersey, USA) and 50 ng/mL IL-6 (Peprotech, New Jersey, USA) throughout the culture period, while 1 ng/mL IL-3 (Peprotech, New Jersey, USA) was included only during the first three weeks.

Mast cell morphology was assessed by Toluidine blue staining. Mast cell purity was determined by flow cytometry, and used when >95% of live cells were FcεRI and CD117 positive. Flow cytometry was performed on LSR Fortessa (BD Bioscience, New Jersey, USA) and data were analyzed using FlowJo v10.6 (FlowJo LLC, Ashland, OR, USA).

### Mast cell degranulation/Beta hexosaminidase assay

Mast cells were identified by CD45+FceRI+CD117+SSC^lo^. To measure mast cell degranulation, we examined levels of membrane CD63, an established marker of mast cell degranulation^11–13^. Mast cell degranulation was also assessed using the β-hexosaminidase assay as previously described^14^. In brief, primary mast cells derived from CD34+ progenitors were stimulated with esophageal lysate for 30 minutes. Following stimulation, secreted and total enzyme activities were measured using supernatants and lysates incubated with p-nitrophenyl N-acetyl-β-D-glucosamide (PNAG) for 90 minutes.

### Quantification of Cytokines

C3 ELISA (Hycult Biotech, Uden, Netherlands) and C3a ELISA (Thermo Fisher Scientific, Waltham, MA) were performed on supernatants of human esophageal biopsy lysates. Esophageal biopsies were sonicated in the presence of protease inhibitor, spun down, and supernatants were used for ELISA. Mast cell supernatant was interrogated for a 96 panel of cytokines (Eve technologies, Calgary, CA). Those which were significant overall were graphed.

### PCR amplification of *C3AR1*, and representative EoE gene analysis

Fresh biopsy specimens were stored in RNAlater until they were subjected to RNA isolation using the miRNeasy kit (Qiagen) per the manufacturer’s instructions. Esophageal biopsy RNA were isolated. After RNA quantity and quality analyses with a NanoDrop spectrometer, an aliquot of 500 ng of RNA was acquired for reverse transcription by using the iScript cDNA Synthesis Kit (Bio-Rad Laboratories), according to the manufacturer’s protocol.

In addition to *C3AR1* transcripts, the transcriptomic signature of esophageal biopsy samples was obtained using an EoE Diagnostic Panel (EDP) comprising a set of 96 esophageal transcripts (which includes housekeeping genes) as previously reported.^9, 10^ *C3AR1* (Hs00269693_s1) and representative EoE genes in EDP were amplified from cDNA stock generated by the methods described above. Using TaqMan Universal Master Mix II (Applied Biosystems), TaqMan real-time PCR amplification was performed on the Quant Studio 7 (Life Technologies). After the qPCR was complete, raw cycle threshold (CT) values for each sample/each gene were exported into GeneSpring GX 12.6 (Agilent Technologies) for statistical analysis. Glyceraldehyde-3-phosphate dehydrogenase (*GAPDH*) was used as an expression control for all analyzed genes. Samples with a *GAPDH* value of <30 CT value were considered acceptable for analysis. The expression CT value of the housekeeping gene *GAPDH* was subtracted from each gene of interest (GOI) CT value to acquire the ΔCT calculated.^9, 10^

### RNA sequencing

Previous bulk RNA sequencing of human esophageal tissue was analyzed using GeneSpring for gene correlations. Previous single cell RNA sequencing data was analyzed for complement specific genes^15^.

### Immunofluorescence

Immunofluorescence was performed as previously described^7^. Briefly, formalin-fixed, paraffin-embedded slides underwent deparaffinization with xylene, followed by graded ethanol washes and antigen retrieval under pressurized conditions. The samples were blocked in 5% donkey serum. Primary antibody staining was performed (anti-C3a; Complement Tech, Texas, USA), with appropriate isotype controls used as negative control and subsequently incubated with fluorescent secondary antibodies (donkey anti-rabbit Alexa Fluor 647). Of note, this is a polyclonal antibody which cannot distinguish between C3a and C3. Sequential, partially overlapping images were captured on a Nikon A1R LUN-V inverted confocal microscope (Nikon Instruments, Melville, NY) at 20× and 40x magnification and stitched to create whole biopsy images. Immunofluorescent images of human esophageal mast cells (anti-tryptase) and fibroblasts (anti-vimentin staining, spindle shape in lamina propria) were obtained from the Enable Medicine platform, and closest distance between border of mast cells and fibroblasts were measured.

### Endoscopic phenotypes

Endoscopic features were prospectively recorded in real-time using a classification and grading system by a simplified endoscopic severity score (ESS) with each feature (edema, rings, exudates, furrows, and strictures) scored as absent or present. The endoscopic phenotype was based on the results of the ESS.^10^ An inflammatory endoscopic phenotype was defined by findings limited to furrows, exudates, or edema, whereas a fibrostenotic endoscopic phenotype was defined if there were findings of rings or esophageal stricture. A normal endoscopic phenotype had no abnormalities visualized on endoscopy.

### Statistical Analyses

Statistical analyses were performed with GraphPad Prism 10.1 (GraphPad Software, San Diego, California). Comparisons between two groups were performed using Mann-Whitney U test given non-normality of data. False discovery rate was accounted for in p value in consideration of significance. For the multiplex data, one way ANOVA was performed for each cytokine, and those with p values ≤0.05 underwent post hoc analysis (Tukey post-test) between individual groups.

## Results

### Mast cell activation in EoE

We hypothesized that there would be increased mast cell degranulation in EoE compared to control. Accordingly, we examined levels of membrane CD63, an established marker of mast cell activation^11–13^ on esophageal mast cells by flow cytometry. Mast cells, identified as CD45+FceRI+CD117+SSC^lo^ (Figure 1A), demonstrated increased surface expression of CD63 in active compared with remission subjects (4.4 fold, p=0.016) and in active compared with control subjects (18.4 fold, p= 0.004) (Figure 1B). We reasoned that mast cells were likely activated in-situ by local factors. Given the lack of benefit of omalizumab in attenuating EoE symptoms^16^, we further hypothesized that this would be by a non-IgE mediated mechanism. To test this hypothesis, esophageal biopsies were lysed, and the supernatant was incubated with primary mast cells which were *not* sensitized with IgE. Compared to buffer, biopsy lysates led to increased degranulation in a dose-dependent manner by flow cytometry (16.9 fold, p=0.0005) (Figure 1C) and by beta hexosaminidase assay (38.8 fold, p=0.01) (Figure 1D).

**Figure 1:**
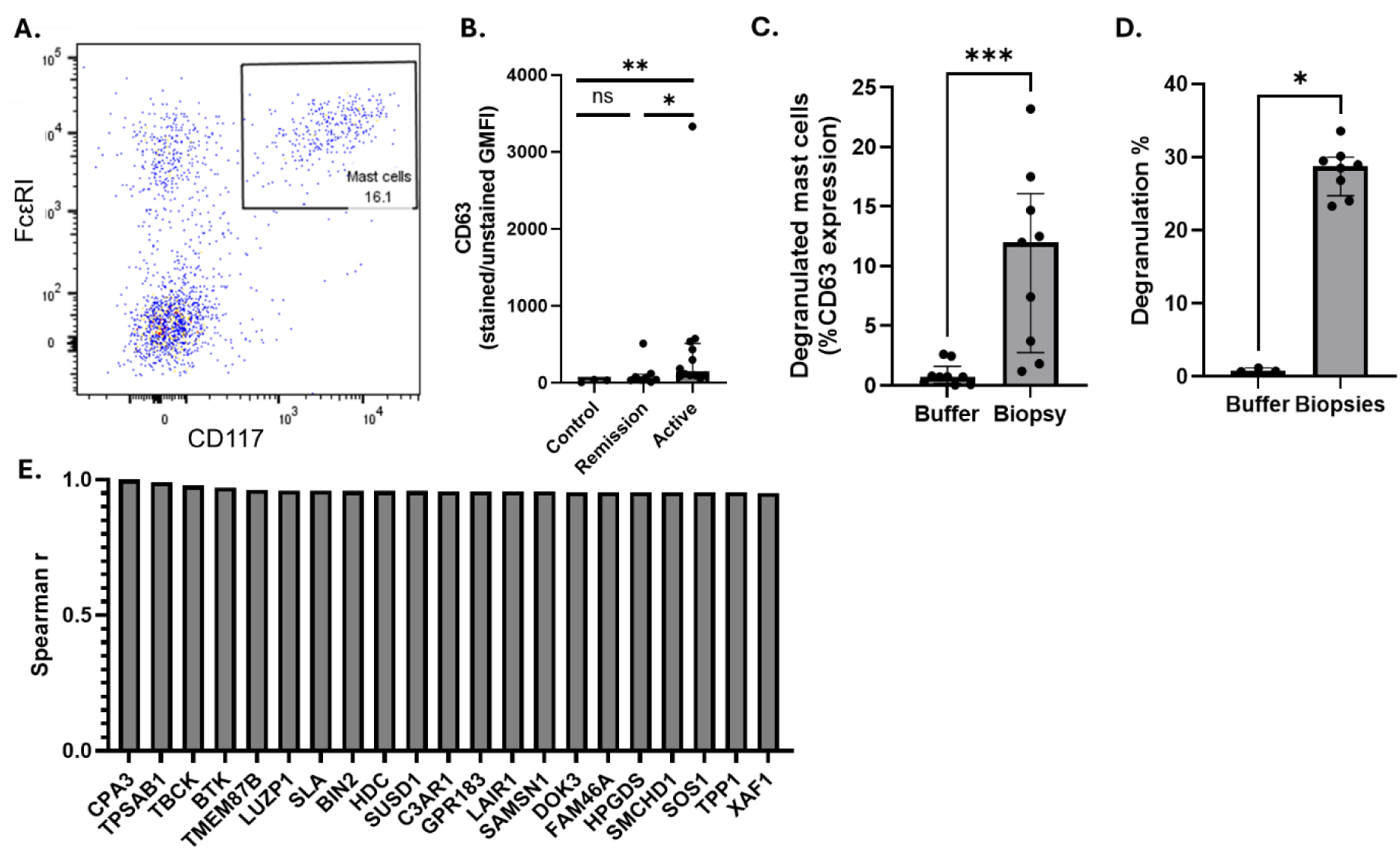
Human esophageal mast cells identification and activation. **A.** A cohort of human esophageal biopsies was analyzed using flow cytometry to identify mast cells (FcERI+ CD117+). The percentage of activated esophageal mast cells, defined by the expression of the activation marker CD63, was compared between control (n=3), remission EoE (n=8), and active EoE (n=12) patients. **B.** The degranulation of mast cells in human esophageal biopsy lysates was assessed using flow cytometry measuring the percentage of CD63 expression and **C.** beta-hexosaminidase assay, with results compared to buffer control. D. Representative genes significantly correlated with the mast cell marker CPA3 were identified through bulk RNA sequencing analysis. Top 20 genes are shown. Mann-Whitney test used to compare between groups. Individual markers represent distinct individuals. Bars represent median with interquartile range. Statistical differences between groups ns=non significant, *p ≤ 0.05, **p < 0.01, ***p < 0.001.

We then aimed to determine potential mechanisms of mast cell activation. We hypothesized that genes which correlated most highly with mast cell specific genes were ligands or receptors which were important in esophageal mast cell activation. Accordingly, bulk RNA sequencing data was probed for genes most highly correlated to the established mast cell marker *CPA3* which is markedly overexpressed in the biopsies of EoE patients^15^. From the EoE transcriptome, the 20 genes which correlated most highly to *CPA3* were *TPSAB1* (Spearman rho= 0.99), *TBCK* (Spearman rho= 0.98), *BTK* (Spearman rho= 0.97), *TMEM87B* (Spearman rho= 0.96), *LUZP1* (Spearman rho= 0.96), *SLA* (Spearman rho= 0.96), *BIN2* (Spearman rho= 0.96), *HDC* (Spearman rho= 0.96), *SUSD1* (Spearman rho= 0.96), *C3AR1* (Spearman rho= 0.96)*, GPR183* (Spearman rho= 0.96)*, LAIR1* (Spearman rho= 0.96)*, SAMSN1* (Spearman rho= 0.96)*, DOK3* (Spearman rho= 0.95)*, FAM46A* (Spearman rho= 0.95)*, HPGDS* (Spearman rho= 0.95)*, SMCHD1* (Spearman rho= 0.95)*, SOS1* (Spearman rho= 0.95)*, TPP1* (Spearman rho= 0.95), and *XAF1* (Spearman rho= 0.95) (Figure 1E).

### C3aR is a mechanism of esophageal mast cell activation

One of the most highly correlated genes to *CPA3* was *C3aR1*, encoding the receptor for C3a (R^2^=0.92, p<0.0001) (Figure 2A). *C3AR1* is also highly correlated with *TPSAB1*, another mast cell specific gene (R^2^=0.82, p<0.0001) (Figure 2B). Flow cytometric analysis demonstrated that the ratio of C3aR GMFI (stained over unstained) was not significantly different between control (n=2), remission EoE (n=8), and active EoE (n=10) patients, although there is a trend towards higher C3aR from remission to active EoE (Figure 2C). There was no difference between the percentage of mast cells that express C3aR between control (n=2), remission EoE (n=8), and active EoE (n=10) patients (Figure 2D). We hypothesized that if C3aR may be involved in mast cell activation, its level of expression might correlate with CD63. This was indeed found (R^2^=0.46, p=0.0007) (Figure 2E).

**Figure 2.**
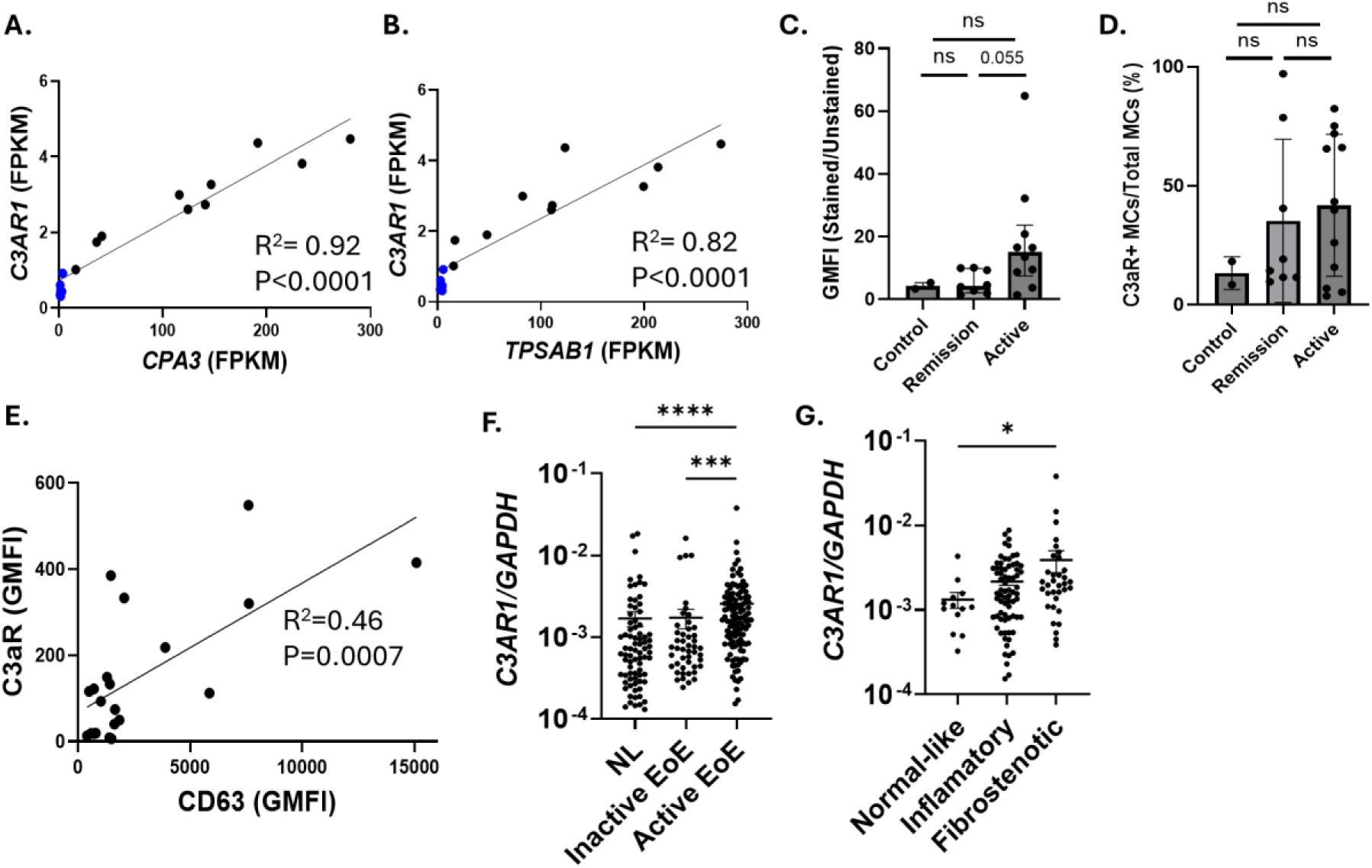
C3a Receptor expression on human esophageal patients. **A, B.** Spearman rank correlation of mast cell specific genes (*CPA3* and *TPSAB1*) with the gene *C3AR1* in subjects with active EoE (n=10) and control (n=6). Simple linear regression was used for correlation. **C**. A cohort of human esophageal mast cells from control (n=2), active (n=10) and remission (n=8) EoE patients was compared to determine the levels of C3aR expression and **D.** the percentage of C3aR+ mast cells (MCs) out of total mast cells. Mann-Whitney test used to compare between groups. Bars represent median with interquartile range. **E.** Correlation of mast cell C3aR with mast cell CD63 on control (n=2), active (n=10) and remission (n=8) patients by simple linear regression. **F.** *C3AR1* transcripts from control (NL; n= 78), remission (inactive EoE; n=47), and active EoE (n=118). **G.** *C3AR1* transcripts from active EoE (n=118) were classified into normal-like, inflammatory, and fibrostenotic phenotypes based on endoscopy. Individual markers represent distinct individuals. Statistical differences between groups ns=non significant.

To examine the potential clinical significance of C3AR1 in EoE, *C3AR1* expression was validated in an independent cohort (118 active EoE, 47 inactive EoE, and 78 controls). Esophageal *C3AR1* expression was markedly higher in patients with active EoE than inactive EoE and controls (Figure 2F), as was the expression of other complement genes, such as *CFB* and *CFI* (Supplementary Figure 1A). Among patients with active EoE, patients with fibrostenotic features had significantly higher *C3AR1* expression than did patients with inflammatory and normal endoscopic features (Fig. 2G). Also, significant correlations were noted between *C3AR1* and the majority of genes involved in EoE (Supplementary Figure 1B).

### Complement dysregulation and activation

We next aimed to determine the expression of complement proteins in the esophagus as a surrogate for complement activation. As an initial inquiry, we examined the bulk and single cell RNA sequencing data from the esophagus of EoE and control individuals for the expression of complement related genes. Multiple genes which were increased in EoE, including *MBL1P, CFI, CFB, C2, ITGAM, CFH, C3AR1, C5AR1, C1QB, C1QA, C1QC, ITGB2-AS1, ITGB2*, and *CD55* (Figure 3A). There were no complement genes which were decreased. Single cell RNA sequencing data showed enrichment of complement genes in multiple cell types, including *C3AR1* in mast cells and *C3* in fibroblasts (Figure 3B). Fibroblasts were enriched for multiple complement genes, including *C1R*, *C1S*, *C2*, *CFB*, *C3*, *C5*, *C7*, *CFD*, *MASP1*, *CFH*, *CD59*, *CFI*, and *SERPING1*. Myeloid cells were also enriched in several complement genes, including *C1QA, C1QB, C1QC, C5, CFP, FCN1, IITGAM, ITGB2, ITGAX, C5AR1*, and *C3AR1*. *CD59* was enriched in multiple cell types, including mast cells, fibroblasts, endothelium, differentiating epithelium, and differentiated epithelium.

**Figure 3.**
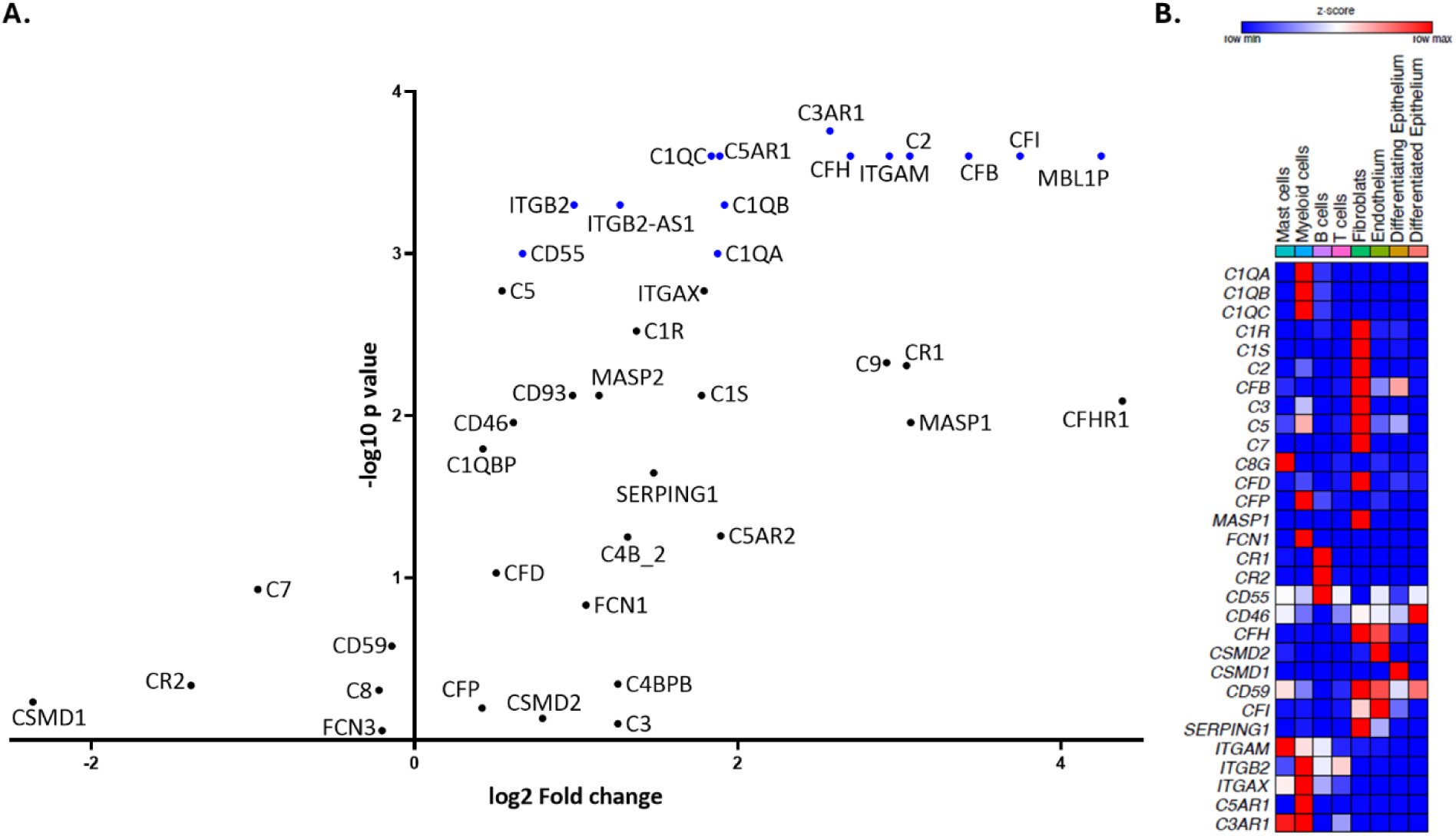
Complement dysregulation in EoE. **A**. Complement genes from esophageal bulk RNA expression. **B.** Heatmap of complement–enriched genes (x-axis) in multiple cell types (y-axis). The map’s color for each gene is proportional to the average per-cell gene expression within the given patient in the esophageal cell populations. Esophageal biopsies derived from patients with active EoE or remission EoE and non-EoE controls.

Given the number of increased compared to decreased complement genes, and that the proteins may undergo a host of post-translational modifications, we next decided to interrogate complement system proteins.

Immunofluorescence showed the presence of immunoreactive C3a (which includes C3) in epithelium and lamina propria (Figure 4A). Furthermore, we hypothesized that there is increased local complement activation in the esophagus in EoE due to an increased number of proteases which may cleave C3 to C3a^17, 18^. Indeed, the ratio of C3a/C3 was increased in EoE compared to control (Figure 4B). There was no difference in plasma C3 between EoE and control, and the ratio of C3 and C3a over total protein were not different between EoE and control (Supplemental Figure 2). Given the enrichment of C3 in fibroblasts, the presence of mast cells in the lamina propria, and the chemotactic ability of C3a on mast cells^19^, we hypothesized that mast cells and fibroblasts are in closer proximity in EoE compared to control. Indeed, this was observed with the median distance decreasing from 2.6 μM in control to 0.2 µM in active EoE biopsies (p=0.01) (Figure 4C).

**Figure 4.**
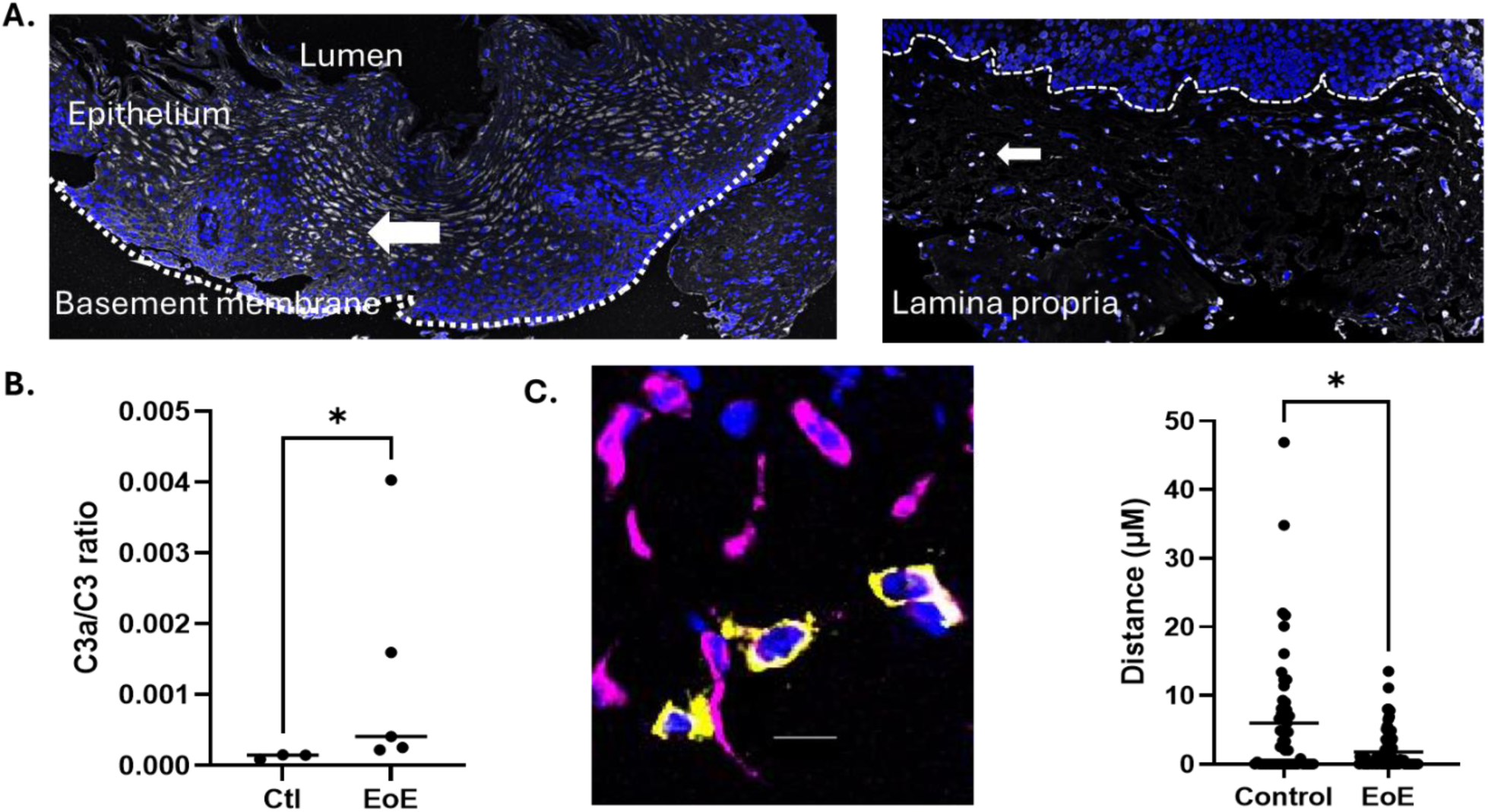
Complement activation in EoE. **A.** Representative pictures of a digitally stitched whole biopsy immunofluorescence of human esophageal tissue, multiple 20× high powered fields. The samples were stained nuclear (blue) and C3a (white). White arrows point towards C3a+ cells. **B.** C3a and C3 from EoE (n=5) and control (n=3) patients. Mann-Whitney test used to compare between groups. Bars represent median with interquartile range. **C.** Immunofluorescent staining of mast cells by anti-tryptase (yellow) and fibroblasts by anti-vimentin (magenta). Distance between each mast cell-fibroblast pair in the esophageal lamina propria in EoE (n=3) and control (n=4) patients. Individual markers represent distinct individuals. Statistical differences between groups *p ≤ 0.05.

### Downstream effects of C3aR stimulation

We aimed to determine the downstream effects of C3aR stimulation of mast cells, with a focus on responses that may be involved in EoE. Primary human mast cells grown from CD34+ progenitors were stimulated with C3a (100 ng/mL) and examined for secretion of a pleiotropy of relevant cytokines kinetically (4 and 8 hours) (Figure 5A). There were several cytokines that were markedly released at 4 hours including granzyme B (7.6-fold, p<0.001), M-CSF (2.7-fold, p=0.01), IL-34 (1.7-fold, p=0.02), and IL-22 (6.2-fold, p=0.01) (Figure 5B). Some remained elevated at 8 hours granzyme B (3.9-fold, p=0.04), M-CSF (2.1-fold, p=0.03), IL-34 (2.4-fold, p<0.001), and IL-22(5.2-fold, p=0.01). Granzyme B was the highest marker on the panel, with an average of 318 pg/mL across all 3 conditions in triplicate. Comparatively, perforin had an average of 2.33 pg/mL.

**Figure 5.**
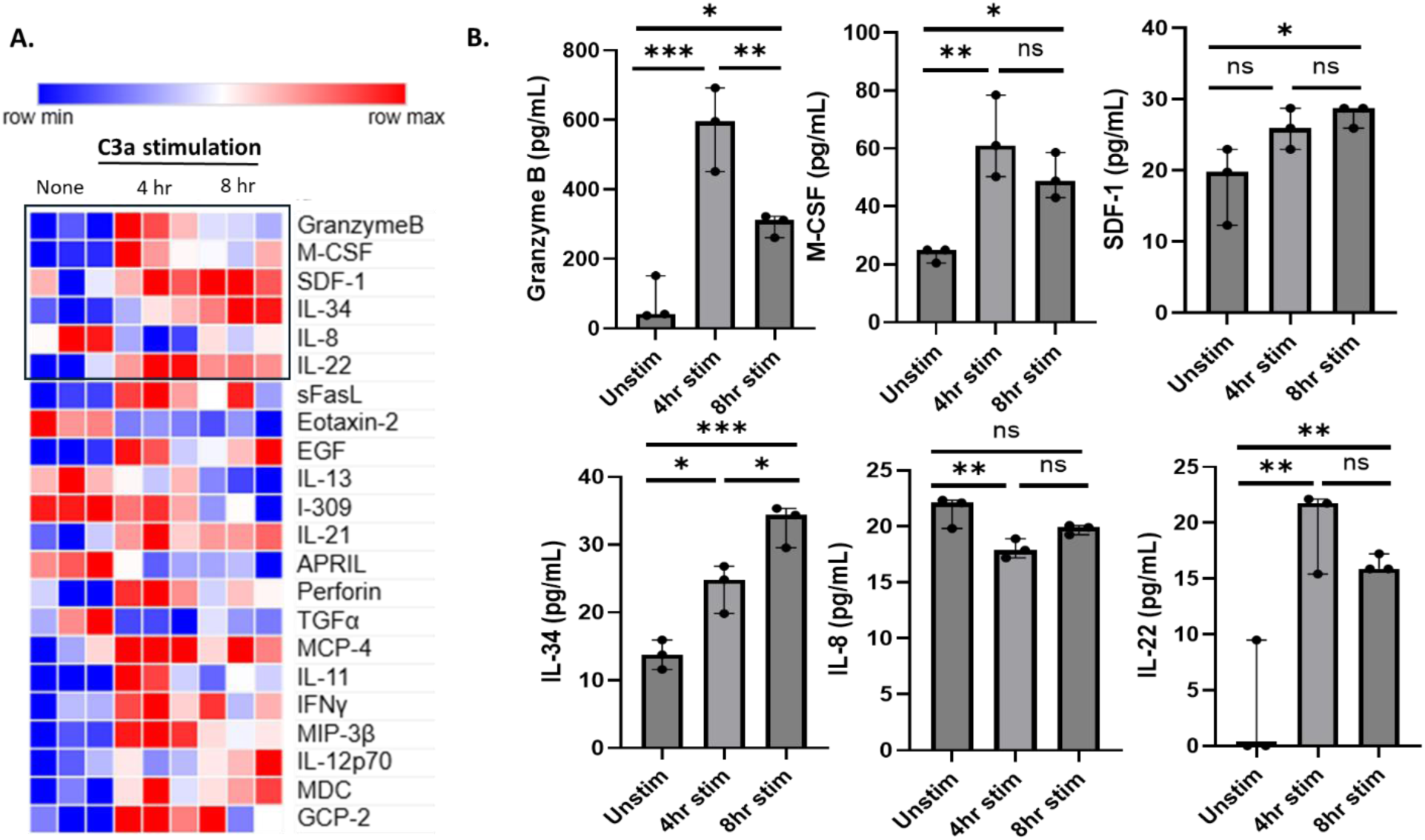
Downstream effects of C3aR stimulation. **A**. Heatmap of the cytokines (y-axis) released by the primary mast cells after the stimulation with buffer (“None”) or 100 ng/ml recombinant C3a in triplicate at 4 and 8 hours (hr) (x-axis). Black box represents all cytokines whose values average more than 10 pg/mL. **B.** Graphs of individual cytokines. Statistical differences between groups *p ≤0.05, **p ≤0.01, ***p ≤0.001, ns = not significant.

### Conclusion

EoE is an allergic inflammatory disease of the esophagus with mast cell involvement and likely activation. In this study, we aimed to determine potential mechanisms of mast cell activation and contribution to disease. We found that 1.) mast cell activation is increased in EoE, potentially through a non-IgE mediated mechanism, 2.) C3a receptor may be involved, 3.) there is increased complement dysregulation in EoE, and that 4.) the interplay of mast cells and fibroblasts may be important in EoE pathogenesis.

Consistent with previous data, esophageal mast cells have increased markers of degranulation in active disease compared to EoE in remission. We have also shown the ability of esophageal lysates to degranulate primary human mast cells as assessed by the release of beta hexosaminidase and flow cytometric analysis for CD63. This provides a valuable tool in further delineating the mechanisms of mast cell activation in the future.

Given the evidence for mast cell activation in EoE, we probed bulk RNA sequencing data to look for genes which correlated most with *CPA3*, the most highly upregulated mast cell specific gene in the EoE transcriptome, and found that *C3AR1* is one of the most correlated genes. This is particularly interesting as C3a is a well-known anaphylatoxin and chemotactic factor for mast cells^19–21^, and we recently also found evidence that suggested mast cell chemotaxis into the epithelium in EoE from the esophageal papillae^7^. Presence of C3a receptor was confirmed on esophageal mast cells, and there is a trend towards increased C3aR on active versus remission mast cells. The C3aR also correlates significantly with surface CD63 expression, which is a measure of mast cell degranulation. This suggests that C3a may be a mechanism for esophageal mast cell degranulation in EoE. Gene expression data with *C3AR1* suggests this may be important in the fibrostenotic phenotype in EoE.

We probed both bulk and scRNA sequencing data for complement dysregulation, and found several complement genes which were increased, including complement factor B (CFB) which is a part of the alternate pathway of complement activation and suggests that increased complement activation in EoE may be via the alternative pathway^22^. There is an increase in regulators of the complement system, including complement factor H (CFH) and complement factor I (CFI) which may be a compensatory mechanism. An increase of complement genes is also seen in other atopic diseases such as asthma and atopic dermatitis^23–25^. In the scRNA sequencing data, many complement proteins are enriched in stromal cells, including fibroblasts and endothelium. This is consistent with the paradigm that local stromal cell inflammation may lead to complement activation which then activates and recruits immune cells like mast cells and myeloid cells^26^. Out of the 8 cell types identified by our scRNA sequencing data, fibroblasts are the most enriched for *C3*. Another independent study showed that fibroblast specific *C3* is increased in EoE compared to control and that the C3 (fibroblast) – C3aR (mast cell) relationship may be important in EoE^8^. *C3* has also been shown to be the most upregulated esophageal gene in an animal model of EoE compared to controls^27^.

There is increased protease activity in EoE, including preformed mast cell mediators like chymase and cathepsin G, which can cleave C3 into C3a^28–31^. Indeed, we found an increased C3a/C3 ratio in local esophageal tissue, which suggests increased local esophageal complement activation. Local complement activation has been observed in other diseases as well^32, 33^. Although *C3* transcripts are primarily enriched in fibroblasts, immunostaining for C3a/C3 shows its presence in both the epithelium and lamina propria, which is consistent with other tissues^32, 34^.

We additionally found increased mast cell and fibroblast proximity in EoE compared to control. Previous research suggested that mast cells from the papillae migrate to the epithelium in active EoE, and our current data suggests that the mast cells in the lamina propria may also migrate, possibly because of C3 from fibroblasts being cleaved to C3a, which then acts as a chemotactic factor for mast cells^7^. It is unclear what serves as the initial C3 convertase which cleaves C3 to C3a-whether it is mast cell proteases released by an alternative method, such as IgE^35^, or whether it is another trigger such as IgG^16, 36^. Given the known role of food in triggering EoE, and the potential of food to trigger complement, it is possible that different foods are contributing to complement activation and local tissue inflammation. Human esophageal fibroblasts stimulated by tumor necrosis factor superfamily member 14 (TNFSF14, also known as LIGHT) increase *C3* as well as other inflammatory mediators ^37^.

C3a stimulation of primary human mast cells resulted in release of granzyme B with comparatively lower amount of perforin. Granzyme B has recently been shown to be increased in EoE biopsies, and this study shows that mast cells may be a source^38^. C3a stimulation resulted in increased M-CSF (CSF-1), SDF-1 (CXCL12), IL-34, IL-8, and IL-22. Transcripts for M-CSF are increased in bulk RNA sequencing of esophageal biopsies^15^. M-CSF has known roles in monocyte/macrophage maturation and survival^39^; M-CSF dependent macrophage recruitment can promote esophageal fibrosis in a recent mouse model^40^.

One limitation of this study is the need to use primary human mast cell lines to study effects of C3a, as tissue resident mast cells respond to the local microenvironment. Another limitation is the lack and variability of lamina propria in human esophageal samples, which can make analysis challenging.

In conclusion, we have further dissected potential mechanisms of mast cell activation in EoE to include a non-IgE mediated mechanism. We have revealed complement dysregulation and increased local activation. We have also revealed potential activating ligand-receptor interactions between mast cells and fibroblasts which may contribute to local inflammation. These findings reveal potential future therapeutic targets.

## Author contributions

S.Z. and M.E.R. conceived the study. S.Z., D.C., and M.E.R. wrote the manuscript. S.Z., D.C., and M.X. conducted experiments with technical and analytical support from J.M.C. and M.R. N.B.B.M. provided analysis of single cell RNA sequencing data. J.D., J.F. and N.B.B.M. provided flow cytometry technical support. T.S. provided analytical support for gene expression data. S.B. and V.A.M obtained human esophageal samples. T.D. provided support in growth of primary human mast cells from CD34+ progenitors.

## Acknowledgements and Sources of Funding

We thank the patients who participated in the research, the staff of the Cincinnati Center of Eosinophilic Diseases, the Research Flow Cytometry Core, the Digestive Diseases Research Core Center and the Bio-Imaging and Analysis Core who made this research possible. We thank Shawna Hottinger for editorial assistance. This work was supported by grants NIH R01 AI045898 and AI124355 and the Campaign Urging Research For Eosinophilic Diseases (CURED) Foundation to M.E.R and Food Allergy Research and Education (FARE) New Investigator Award (to S.Z.).

---

**Supplemental Figure 1.**
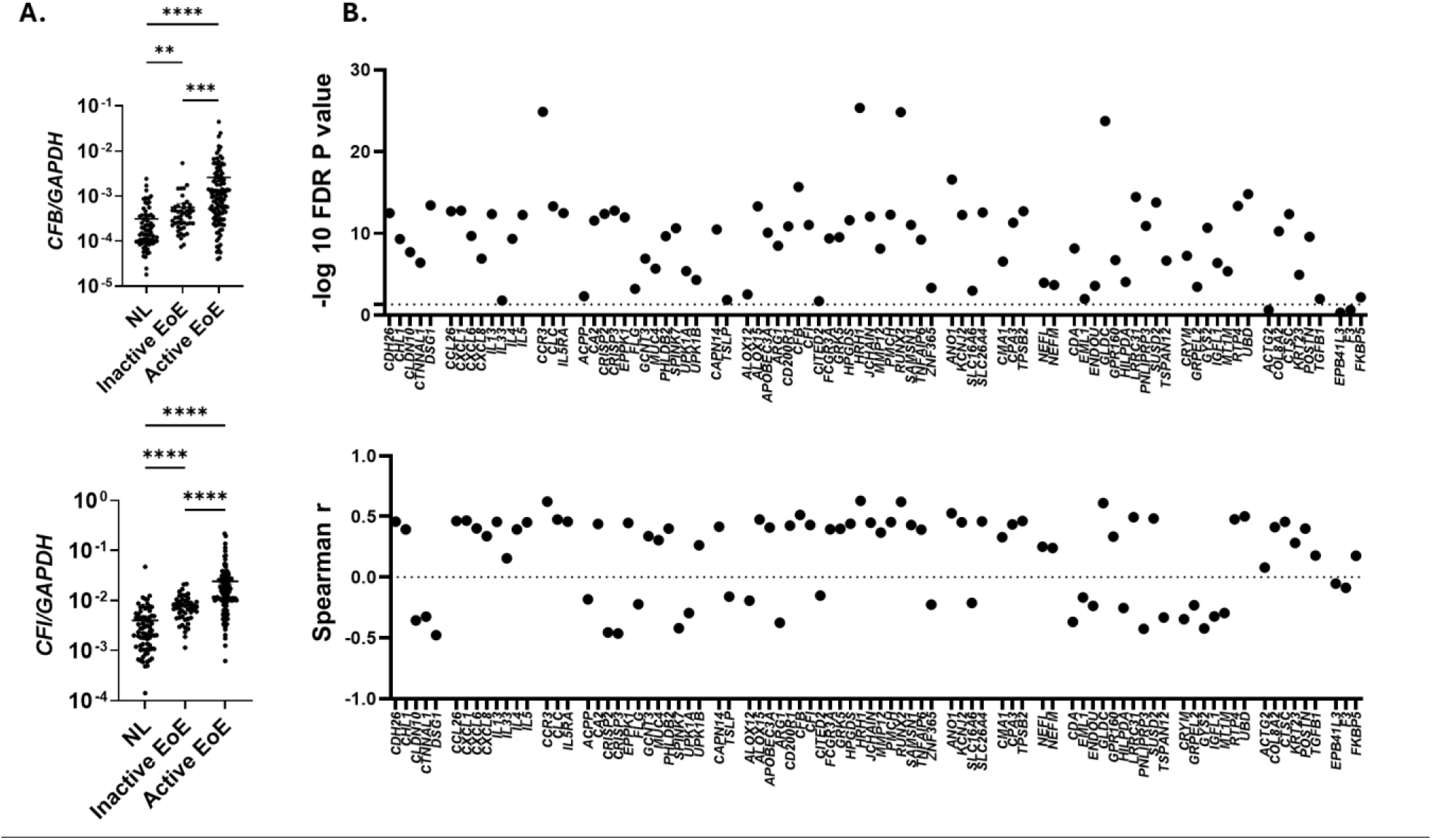
**A.** *CFB* and *CFI* transcripts from control (NL; n= 78), remission (inactive EoE; n=47), and active EoE (n=118). **B.** P value and Spearman r correlations between *C3AR1* and Eosinophil Diagnostic Panel genes.

**Supplementary Figure 2.**
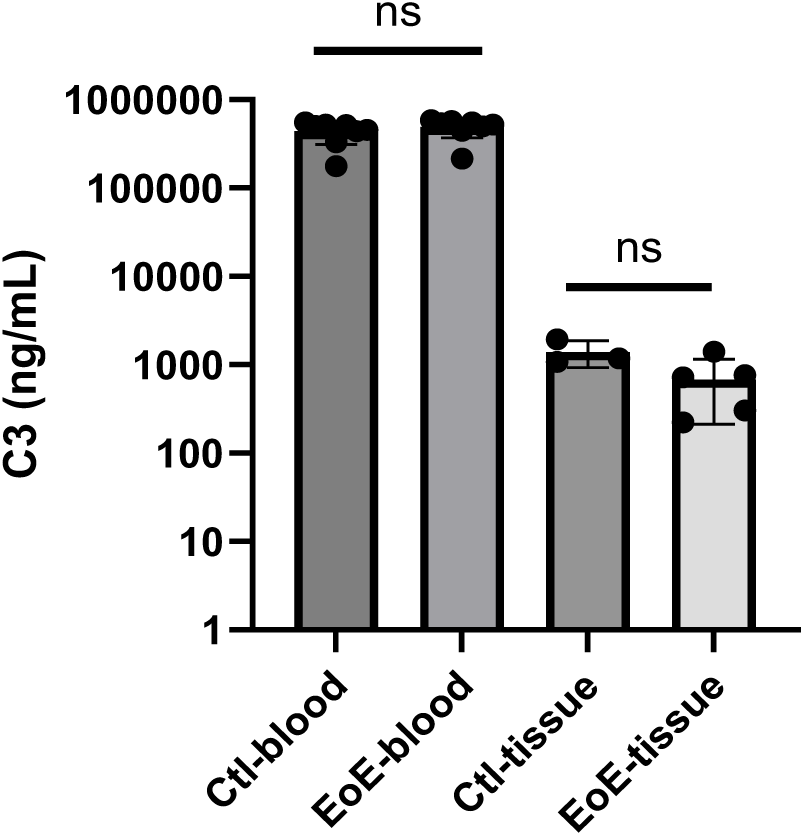
C3 in control (blood-n=8, tissue n=3) and EoE (blood-n=8, tissue n=5) patients. NS=non significant.

